# A *cis*-regulatory variant in *ASIP* causes gray coat color in the donkey

**DOI:** 10.64898/2026.06.23.733953

**Authors:** Yuanyuan Li, Yu Liu, Jiahui Wu, Shuqin Liu, Xiaoran Lin, Kaibo Guo, Tao Yang, Mo Feng, Hetong Zhang, Xinyu Wang, Wenhui Xing, Shiyu Qian, Ran Yang, Chunjiang Zhao

**Affiliations:** College of Animal Science and Technology, China Agricultural University, Beijing 100193, China; State Key Laboratory of Animal Biotech Breeding, Beijing 100193, China; Equine Center, China Agricultural University, Beijing 100193, China; College of Animal Science and Technology, Qingdao Agricultural University, Qingdao 266109, China; Beijing General Station of Animal Husbandry, Beijing 100107, China; Department of Management and Engineering, University of Padua, Padova 35122, Italy; National Engineering Laboratory for Animal Breeding, Beijing 100193, China; Key Laboratory of Animal Genetics, Breeding and Reproduction, Ministry of Agriculture and Rural Affairs, Beijing 100193, China; Beijing Key Laboratory of Animal Genetic Improvement, Beijing 100193, China

**Author notes:** Correspondence: Chunjiang Zhao.

## Abstract

**Background:** Gray is one of the relatively rare coat colors in donkeys. The Hetian Gray donkey is a distinctive indigenous breed from the Xinjiang Uygur Autonomous Region of Northwestern China, characterized by progressive hair depigmentation with aging while retaining dark skin pigmentation. However, the genetic basis underlying this unique gray coat color phenotype remains unclear.

**Results:** To elucidate the genetic basis, we conducted whole-genome resequencing of Gray and non-Gray donkeys. Genome-wide selection signature analyses identified a candidate region on chromosome 15. Subsequent fine-mapping using mass spectrometry–based genotyping of 42 loci refined the candidate interval and revealed a SNP within intron 2 of the *ASIP* gene, located in a genomic fragment with highly similar sequences, showing complete association with the gray coat color. Association analysis in an expanded population further confirmed a strong correlation between this variant and the gray phenotype. Gene expression analyses also supported the role of *ASIP* in regulating pigmentation in donkeys.

**Conclusions:** These findings identify a genetic determinant of gray coat color in donkeys and provide new insights into the molecular mechanisms underlying age-related depigmentation in domestic animals.

## Background

Gray coat color shows distinct patterns across different species. In mammals such as rats, dogs, and sheep and rabbits, this gray pigmentation is present at birth and remains remarkably stable throughout their lifetime [1–4]. In contrast, in horses, donkeys, and cattle, gray animals gradually lose hair pigmentation with advancing age [5–8]. The Gray phenotype is regarded as a feature of some breeds and reflects the selection of human beings for favorite coat color, and has a link with human culture, and it is associated with the high incidence of melanoma in the horse [5].

Previous study in horses have identified a causative mutation for gray coat color, consisting of a 4.6-kb duplication within intron 6 of *STX17* [5]. Although some progresses have made in the study on the genetic mechanism of donkey coat color phenotypes, which discovered that the variants of *ASIP*, *TBX3*, and *TYR* are associated with dark coat colors, non-rabbit-brown, color dilution in donkeys, respectively [9–13], the causative mutation underlying the gray coat color has not been identified yet.

Though gray individuals could be found in most indigenous donkey breeds, they usually account for very little proportion in the populations [14]. However, the Hetian Gray donkey, an indigenous breed originally named Guala or Gulo, mainly distributed in Pishan County of Hetian City, Xinjiang Uygur Autonomous Region, China [15], features large stature and gray coat color gradually lightening with aging while maintaining dark skin pigmentation [6, 16, 17]. These features of the Hetian Gray donkey make it a unique genetic resource in Xinjiang, and an ideal population for the genetic study of donkey gray coat color.

Advances in high-throughput, cost-effective sequencing technologies have enabled large-scale genomic studies in livestock. Whole-genome resequencing of individuals from different phenotypic groups allows rapid identification of genomic regions and candidate genes associated with complex and monogenic traits through genome-wide association studies (GWAS) and selection signal analyses [18–22]. In this study, we performed whole-genome resequencing for Hetian Gray donkeys and non-Gray donkeys from several indigenous breeds, and systematically investigated the genetic basis of the gray coat color phenotype by integrating genome-wide selection signature analyses, population validation, and transcript analysis. The findings of our study provide new insights into the genetic architecture of gray coat color in donkeys and broaden the understanding the mechanism of the common gray coat color phenotype in different species caused by distinct genes.

## Methods

### Sample collection

Blood samples of totally 475 unrelated individuals were collected from nine donkey populations, including 90 Dezhou donkeys sampled from Dong’ e county of Shandong province; 30 Guangling donkeys from Guangling county, Shanxi province; 30 Guangling donkeys from a donkey farm from Guangzong county, Hebei province; 35 Guanzhong donkeys from Fufeng county, Shaanxi province; 26 Yunnan donkeys from Chuxiong city, Yunnan Province; 10 Huaibei Gray donkeys from Huaibei City, Anhui province; 7 Tibetan donkeys from Rikaze county, Tibet province; 1 Taihang donkey from the Science and Technology Backyard of China Agricultural University in Yanqing district, Beijing; 15 Xinjiang donkeys and 122 Hetian Gray donkeys from Pishan county, Xinjiang Uygur Autonomous Region.

Except for samples used for first-generation sequencing validation and mass spectrometry genotyping, all remaining individuals were sequenced using whole-genome sequencing (WGS). Whole-genome resequencing data for the Guangling and Dezhou populations have been deposited in GenBank under BioProject accession number PRJNA800192, while data for the Yunnan, Huaibei Gray, and Hetian Gray donkey populations are available under BioProject accession number PRJNA1215270.

Furthermore, 109 publicly available whole-genome resequencing datasets were included in the analysis. These publicly available data were retrieved from the NCBI database under BioProject accession numbers PRJNA675210, PRJNA431818, and PRJNA896558.

### Whole-genome sequencing and variant calling

Genomic DNA was extracted from blood samples using the Rapid Blood Genomic DNA Extraction Kit (Tiangen Technology Co., Ltd., Beijing, China) and randomly fragmented to construct paired-end libraries with an average insert size of 350 bp. Libraries were sequenced on the Illumina HiSeq X platform, generating 150-bp paired-end reads at an average depth of ∼5×. The donkey reference genome (GCA_016077325.1) [12] was obtained from NCBI (https://www.ensembl.org/) and applied in the subsequent data processing.

Raw reads were processed with Trimmomatic v0.39 [23] for quality control and subsequently aligned to the reference genome using BWA v0.7.17 [24, 25]. Alignments were filtered with Samtools v1.10 [26], and variants were called and genotyped using GATK v4.1.7.0 [27] with HaplotypeCaller, SelectVariants, and VariantFiltration. Low-confidence variants were removed based on established thresholds for sequencing depth (QD < 2.0), mapping quality (MQ < 40.0), strand bias (FS > 60.0, SOR > 3.0), and rank sum tests (MQRankSum < −12.5, ReadPosRankSum < −8.0).

SNPs were further filtered in PLINK v1.9 [28] to exclude variants with call rates < 85%, Hardy–Weinberg equilibrium p-values < 1 × 10⁻⁷, missing genotypes > 10%, or minor allele frequency < 5%. Individuals with > 10% missing genotype data were also excluded. After the quality control, 1,435,541 high-confidence SNPs were retained for downstream analyses.

### Mass spectrometry–based genotyping and validation in an expanded population

Genomic DNA was extracted from blood samples of 180 individuals and adjusted to 10–30 ng/μL after quality assessment by agarose gel electrophoresis. Forty-two candidate SNPs were selected for genotyping. PCR and single-base extension primers were designed using Assay Design Suite v3.1 (Agena Bioscience) and purified by PAGE. Target regions (∼120 bp) were amplified in 5 μL multiplex PCR reactions, treated with shrimp alkaline phosphatase, and subjected to single-base extension using iPLEX Pro reagents. Extension products were desalted, crystallized on sample plates, and analyzed by MALDI-TOF-MS with a MassARRAY Compact Analyzer. Genotypes were called and visualized using Typer 4.0 software. SNPs with poor clustering or low call rates were excluded [29–31]. All 42 SNPs were successfully genotyped in 180 samples with an overall call rate of 98.5% [29–31].

### Validation in an extended population

For samples with genotypes derived from resequencing data, raw sequencing reads were subjected to quality control using Trimmomatic v0.39 [23] and subsequently aligned to the reference genome with BWA v0.7.17 [24, 25]. All remaining samples were amplified using primers *ASIP*_long_F and *ASIP*_long_R and KOD One™ PCR Master Mix (Blue), and the resulting PCR products were subjected to Sanger sequencing with internal primers at the Beijing Novogene Genomics Institute (Table S1). Intronic sequences from multiple mammalian species were aligned using MAFFT v7.525 [32]. The alignment of the sequencing results was manually inspected and visualized using Jalview [33].

### RNA extraction and qPCR

Skin tissue samples were collected from a total of 9 donkeys, including three Hetian Gray donkeys with the homozygous genotype, three Hetian Gray donkeys with the heterozygous genotype at the locus of *ASIP*_IN2 and three Sanfen individuals of Dezhou donkeys. For each individual, skin tissues were obtained from both the abdominal and dorsal regions. All samples were immediately preserved in RNA stabilization solution and stored at −80 °C until further analysis.

Total RNA was extracted from skin tissues using TRIzol reagent (Thermo Fisher Scientific, USA) in accordance with the manufacturer’s instructions. RNA concentration and purity were determined using a NanoDrop 2000 spectrophotometer (Thermo Fisher Scientific, USA). First-strand cDNA was synthesized from 1 μg of total RNA using a reverse transcription kit (Tiangen Technology Co., Ltd., Beijing, China).

qPCR was carried out using SYBR Green Master Mix (Accurate Biology, China) on an Accurate Biology real-time PCR system. The thermal cycling conditions were as follows: initial denaturation at 95 °C for 10 min, followed by 40 cycles of denaturation at 95 °C for 15 s and annealing/extension at 60 °C for 60 s. Gene expression levels were normalized with the internal reference gene β-actin, and relative expression levels were calculated using the 2^−ΔΔCt^ method. All experiments were performed in triplicate. Statistical analyses were performed using t-tests or one-way ANOVA as appropriate, with p < 0.05 considered statistically significant.

## Results

### Genome-wide selection signal analysis for candidate region on the genome

Whole-genome resequencing was performed on 30 Gray animals from the Hetian Gray donkey and 60 non-Gray donkeys from Guangling and Dezhou breeds, with an average sequencing depth of approximately 5×, respectively. Genome-wide F_ST_ scans revealed multiple regions exhibiting pronounced differentiation between Gray and non-Gray populations. Notably, a strong selection signal was detected on chromosome 15, spanning 22,734,659–27,603,538 bp, corresponding to the top 1% of the F_ST_ distribution (Fig. 1).

**Fig. 1.**
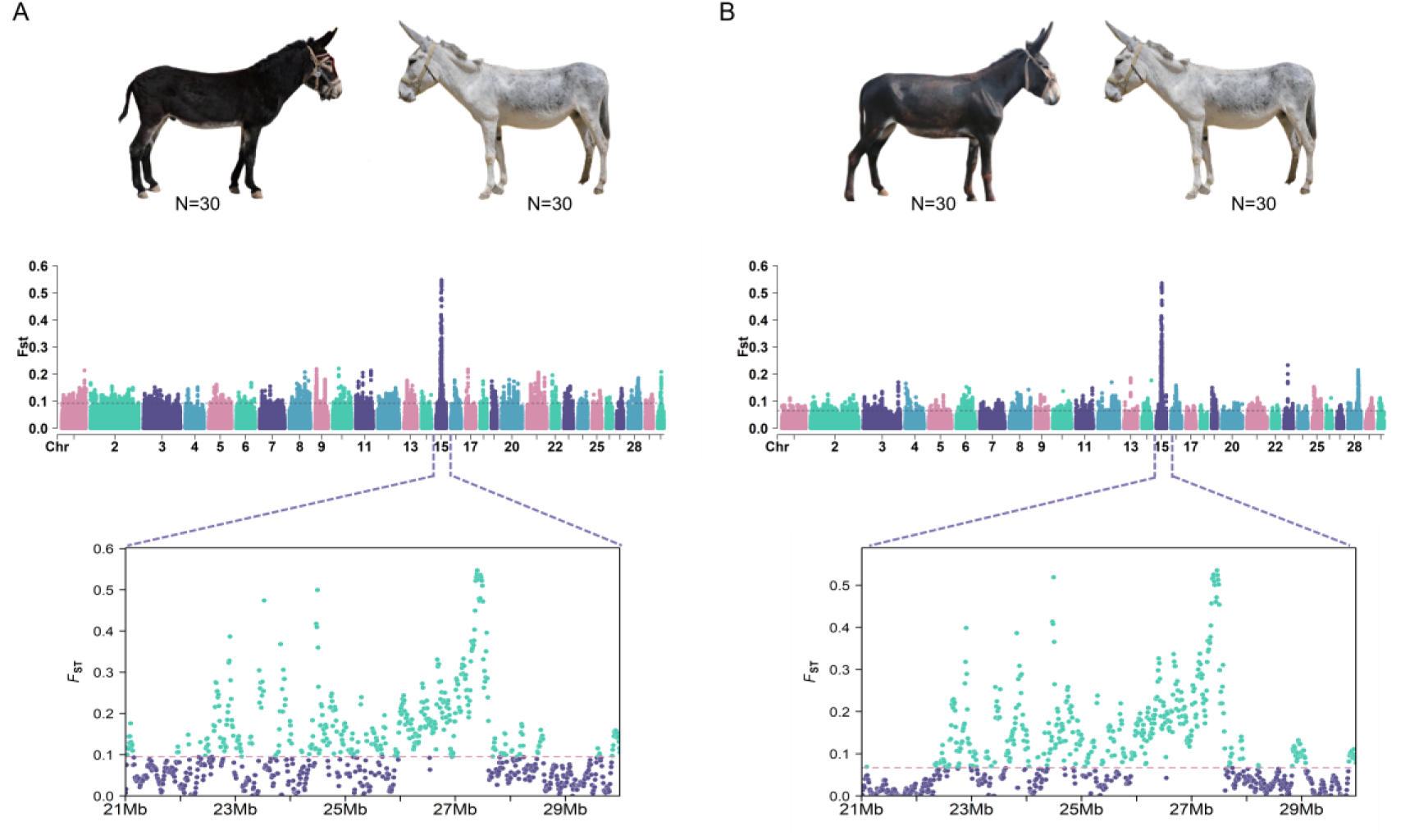
Genome-wide detection of selection signatures between Gray and non-Gray donkeys. **A**: Selection signature analysis and strongly selected genomic regions between Hetian Gray donkeys and Guangling donkeys. **B**: Selection signature analysis and strongly selected genomic regions between Hetian Gray donkeys and Sanfen individuals of the Dezhou donkey. N, Number of the studied donkeys. The dashed line represents the 1% F_ST_ threshold. Images of Guangling donkey, Hetian Gray donkey, and Sanfen individual of the Dezhou donkey are from Animal Genetic Resources in China Horses Donkeys Camels [14].

### Refinement of candidate regions by mass spectrometry-based genotyping

Using a mass spectrometry–based technique, genotyping of 42 single-nucleotide polymorphisms (SNPs) of the preliminary candidate regions with elevated allele frequencies in gray population was performed in 90 Gray donkeys from the Hetian Gray donkey population and 90 non-Gray donkeys representing three breeds (Table S2). Allele frequency comparisons showed that most SNPs within the candidate interval exhibited uniformly high frequencies in Gray donkeys, limiting the resolution achievable based on allele frequency differentiation alone (Fig. 2A).

**Fig. 2.**
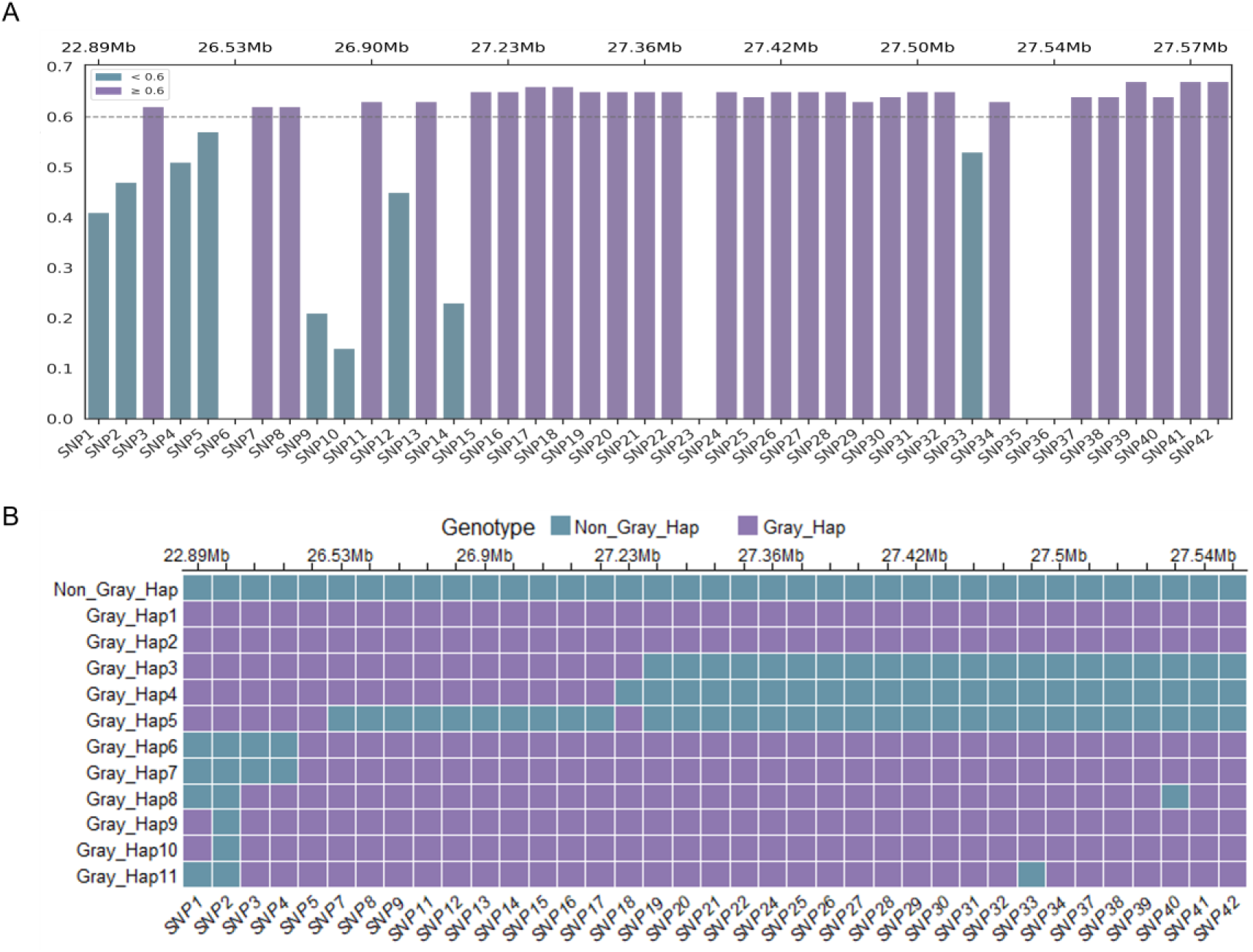
Mass spectrometry–based genotyping results. **A**: The absolute values of allele frequency differences between Gray and non-Gray donkey populations. **B**: Haplotype comparison between Gray and non-Gray donkeys. SNP1-42, the SNPs in the most significantly selected region used for genotyping the Gray and non-Gray donkeys. Non-Gray_Hap, haplotype of non-Gray donkeys; Gray_Hap1 to Gray_Hap11, haplotypes of Gray donkeys.

Further examination of SNPs at which Gray donkeys shared genotypes identical to those observed in non-Gray individuals revealed distinct patterns of marker distribution and linkage across the region. Integrating evidence from allele frequency profiles and genotype-sharing patterns enabled the refinement of the original candidate interval into three discrete genomic sub-regions located at 22,925,355–26,526,995 bp, 27,195,425–27,295,679 bp, and 27,499,274–27,535,127 bp (Fig. 2B). These regions were considered to contain candidate Gray-causing mutations.

### Identification of a variant in *ASIP* completely associated with gray coat color

Based on the mass spectrometry–based genotyping, the three refined genomic intervals were subsequently interrogated using whole-genome resequencing data. Variant calling across these finely mapped regions identified a total of 42 SNPs highly associated with the gray coat color phenotype (Table S3). To prioritize candidate causal variants, all of the 42 SNPs were manually inspected using the Integrative Genomics Viewer (IGV) [34, 35] by comparing aligned sequencing reads (BAM files) from Gray and non-Gray donkeys, with particular emphasis on variants showing consistent presence or increased allele frequency in Gray individuals. Among these candidates, a SNP (A/G), located at the position 26,236,519 bp in the latest reference genome (GCA_041296235.2), exhibited complete association with the gray coat color phenotype. This variant is located within intron 2 of *ASIP* (XM_044748288), a key gene involved in pigmentation, and was therefore designated *ASIP*_IN2. Based on the strong genetic association identified at the *ASIP* locus, we examined the evolutionary conservation of the candidate mutation site to assess its potential functional relevance. The flanking sequences of the mutation were retrieved and aligned with orthologous regions from multiple mammalian species (Fig. S1). This analysis revealed that the candidate site is highly conserved across species, suggesting that it may play an important functional role in gene regulation or pigmentation processes.

This variant could not be reliably genotyped using the mass spectrometry–based assay, likely due to the presence of highly similar sequences in the genome, which interfered with accurate genotype calling (Fig. 3A). To overcome this limitation and ensure target specificity, a 3,272 bp genomic fragment encompassing the candidate SNP was amplified by designing a primer anchoring the specific sequence of the targeted region (Fig. 3B). After the confirmation of single-target amplification, the genotype at this locus was determined by Sanger sequencing using internal primers (Fig. 3B, Table S1), resulting in accurate and reliable genotyping (Fig. 3C).

**Fig. 3.**
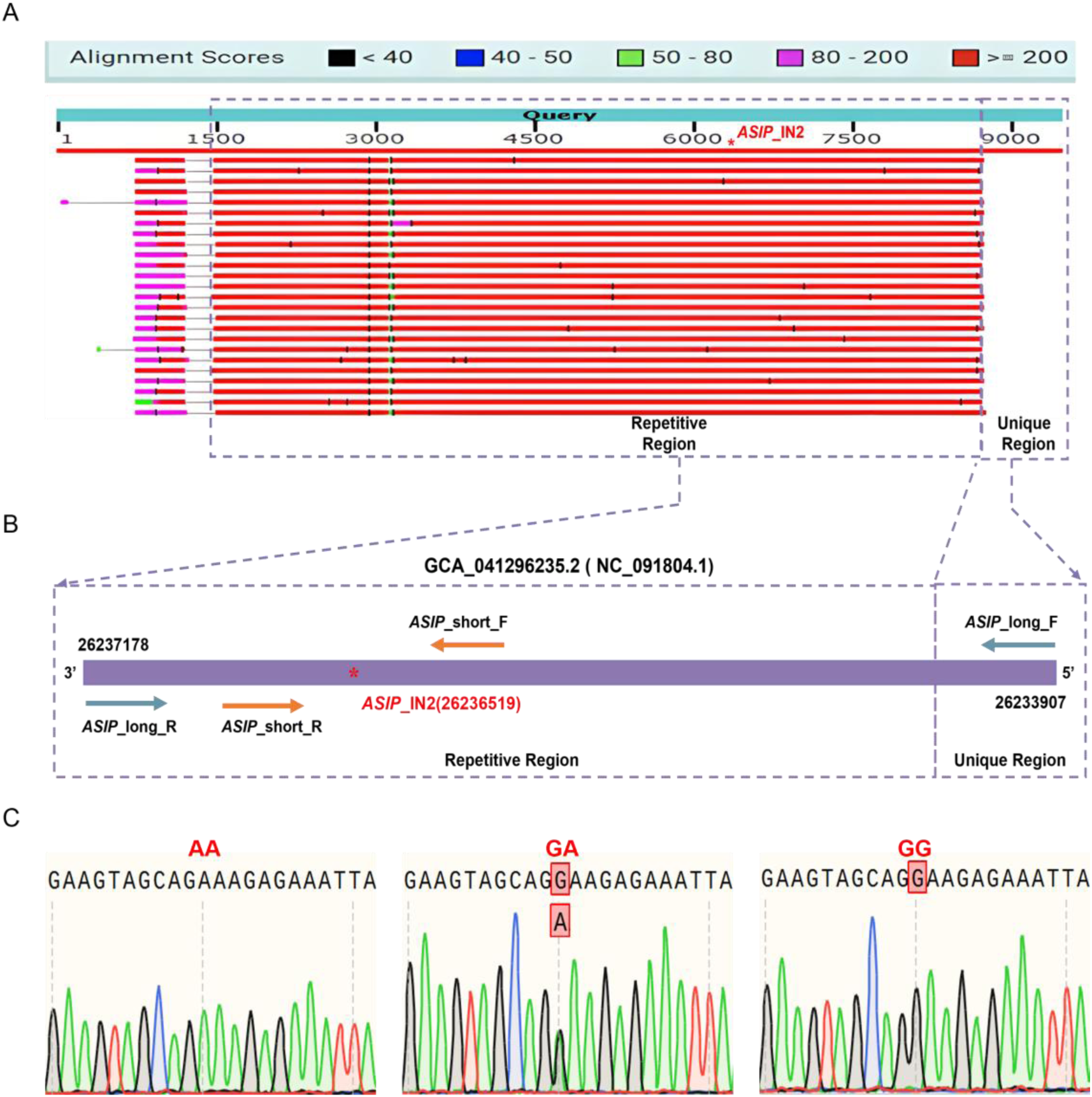
Accurate genotyping of the candidate Gray-causing mutation. **A**: NCBI BLAST alignment results of sequences flanking the candidate variant based on the latest reference genome (GCA_041296235.2), illustrating the highly similar sequences overlapping the targeted region. The repetitive lines indicate the homologous segments that share high sequence similarity on the genome, whereas the unique region denotes a locus-specific sequence. **B**: Schematic representation of the candidate SNP locus based on the latest reference genome (GCA_041296235.2), showing the position of the mutation and the primer design strategy for long-range PCR amplification and internal Sanger sequencing. *ASIP*_long_F, forward primer for the long-range PCR, anchoring the specific sequence of the unique region; *ASIP*_long_R, reverse primer for the long-range PCR; *ASIP*_short_F and *ASIP*_short_R, primers for sequencing the products of the long-rang PCR. **C**: Representative Sanger sequencing chromatograms illustrating the three genotypes (GG, GA, and AA) detected at the candidate SNP locus in Gray and non-Gray donkey populations.

The SNP was validated in a population with well-characterized coat colors, comprising 126 Gray donkeys (122 Hetian Gray donkey, 3 Tibetan donkey, and 1 Taihang donkey) and 70 non-Gray donkeys (20 Dezhou donkey, 20 Guangling donkey, and 30 Guanzhong donkey). G allele of *ASIP*_IN2 was detected in all Gray donkeys but absent in non-Gray donkeys, which harbored a uniform AA genotype (Table 1). The results showed a complete association between the Gray phenotype and *ASIP*_IN2 across donkey breeds.

**Table 1.** Complete association between the Gray phenotype and the mutation in *ASIP* intron 2 across donkey breeds*

| Coat Color | Breeds | N | Genotype |  |  | Genotype frequency |  |  | Allele frequency |  |
| --- | --- | --- | --- | --- | --- | --- | --- | --- | --- | --- |
|  |  |  | GG | GA | AA | GG | GA | AA | G | A |
| Gray | HT | 122 | 37 | 85 | 0 | 0.302 | 0.698 | 0.000 | 0.651 | 0.349 |
|  | TI | 3 | 1 | 2 | 0 |  |  |  |  |  |
|  | TH | 1 | 0 | 1 | 0 |  |  |  |  |  |
|  | Total | 126 | 38 | 88 | 0 |  |  |  |  |  |
| Non_Gray | DZ | 20 | 0 | 0 | 20 | 0.000 | 0.000 | 1.000 | 0.000 | 1.000 |
|  | GL | 20 | 0 | 0 | 20 |  |  |  |  |  |
|  | GZ | 30 | 0 | 0 | 30 |  |  |  |  |  |
|  | Total | 70 | 0 | 0 | 70 |  |  |  |  |  |

\*: HT, Hetian Gray donkey; TI, Tibetan donkey; TH, Taihang donkey; DZ, Dezhou donkey; GL, Guangling donkey; GZ, Guanzhong donkey; N, Number of the studied donkeys.

To investigate the frequencies of G allele at *ASIP*_IN2 locus in diverse donkey breeds except Hetian Gray donkey, we analyzed publicly available whole-genome sequencing data of 189 donkey, encompassing 13 donkey breeds lacking associated individual coat color records, revealed 2 GG, 24 GA, and 163 AA genotypes at the target locus in the populations. The 24 individuals with GA genotype were from Dezhou, Guangling, Guanzhong, Yunnan, Huaibei, Xinjiang, Tibetan, Sichuan, and Qinghai donkeys; GG genotype was detected for one Qinghai donkey and one Shanbei donkey, respectively (Table 2). Overall, in these populations without recorded individual phenotypes, the A allele was predominant (92.6%), whereas the G allele was rare (7.4%), and the results were in accordant with the occurrence of gray coat color in the studied populations investigated previously [14]. These findings further support the strong association between *ASIP*_IN2 and the gray coat color phenotype, and indicate that the G allele occurs only sporadically in indigenous donkey populations except the Hetian Gray donkey, reflecting limited selection for gray coat color in these breeds.

**Table 2.** Genotype and allele frequencies at the target locus in *ASIP* intron 2 across 13 donkey breeds*

| Breeds | N | Genotype |  |  | Genotype frequency |  |  | Allele frequency |  |
| --- | --- | --- | --- | --- | --- | --- | --- | --- | --- |
|  |  | GG | GA | AA | GG | GA | AA | G | A |
| DZ | 33 | 0 | 3 | 30 | 0.000 | 0.091 | 0.909 | 0.046 | 0.955 |
| GL | 10 | 0 | 1 | 9 | 0.000 | 0.100 | 0.900 | 0.050 | 0.950 |
| GZ | 16 | 0 | 1 | 15 | 0.000 | 0.063 | 0.938 | 0.031 | 0.969 |
| YN | 26 | 0 | 3 | 23 | 0.000 | 0.115 | 0.885 | 0.058 | 0.942 |
| HBH | 16 | 0 | 4 | 12 | 0.000 | 0.250 | 0.750 | 0.125 | 0.875 |
| XJ | 20 | 0 | 2 | 18 | 0.000 | 0.100 | 0.900 | 0.050 | 0.950 |
| TI | 10 | 0 | 3 | 7 | 0.000 | 0.300 | 0.700 | 0.150 | 0.850 |
| BY | 6 | 0 | 0 | 6 | 0.000 | 0.000 | 1.000 | 0.000 | 1.000 |
| SCH | 26 | 0 | 1 | 25 | 0.000 | 0.039 | 0.962 | 0.019 | 0.981 |
| QH | 8 | 1 | 6 | 1 | 0.125 | 0.750 | 0.125 | 0.500 | 0.500 |
| QY | 5 | 0 | 0 | 5 | 0.000 | 0.000 | 1.000 | 0.000 | 1.000 |
| TH | 5 | 0 | 0 | 5 | 0.000 | 0.000 | 1.000 | 0.000 | 1.000 |
| SHB | 8 | 1 | 0 | 7 | 0.125 | 0.000 | 0.875 | 0.125 | 0.875 |
| Total | 189 | 2 | 24 | 163 | 0.011 | 0.127 | 0.862 | 0.074 | 0.926 |

\*: DZ, Dezhou donkey; GL, Guangling donkey; GZ, Guanzhong donkey; YN, Yunnan donkey; HBH, Huaibei Gray donkey; XJ, Xinjiang donkey; TI, Tibetan donkey; BY, Biyang donkey; SCH, Sichuan
donkey; QH, Qinghai donkey; QY, Qingyang donkey; TH, Taihang donkey; SHB, Shanbei donkey; N, sample number of the donkeys. The coat colors of the studied donkeys were not recorded.

### Expression of *ASIP* and *MC1R*

Quantitative real-time PCR (qPCR) was performed to assess *ASIP* expression in the heart, liver, skin (dorsal and ventral), spleen, and muscle tissues of homozygous and heterozygous Gray donkeys. According to the latest reference genome (GCF_041296235.1), the *ASIP* gene contains two major transcript types, including a long transcript and a short transcript. Total *ASIP* transcripts were detected in all examined tissues for both genotypes, and the overall expression trends across tissues were consistent between dorsal and abdominal skin (Fig. 4A and Fig. 4B). *ASIP* expression in skin tissues was lower than in the heart and liver but higher than in the spleen and muscle. Given that skin is a key tissue for pigmentation, qPCR analyses were performed on dorsal and abdominal skin tissues from Gray donkeys (the Hetian Gray donkey) and non-Gray donkeys (Sanfen individuals of the Dezhou donkey). Overall, total *ASIP* transcript expression in Gray donkey skin was significantly higher than in non-Gray donkeys (P<0.05). Within Gray donkeys, *ASIP* expression exhibited both genotype- and region-dependent patterns: heterozygous individuals displayed slightly higher *ASIP* levels in abdominal skin, whereas homozygous individuals showed higher expression in dorsal skin (Fig. 4C, Fig. 4D).

**Fig. 4.**
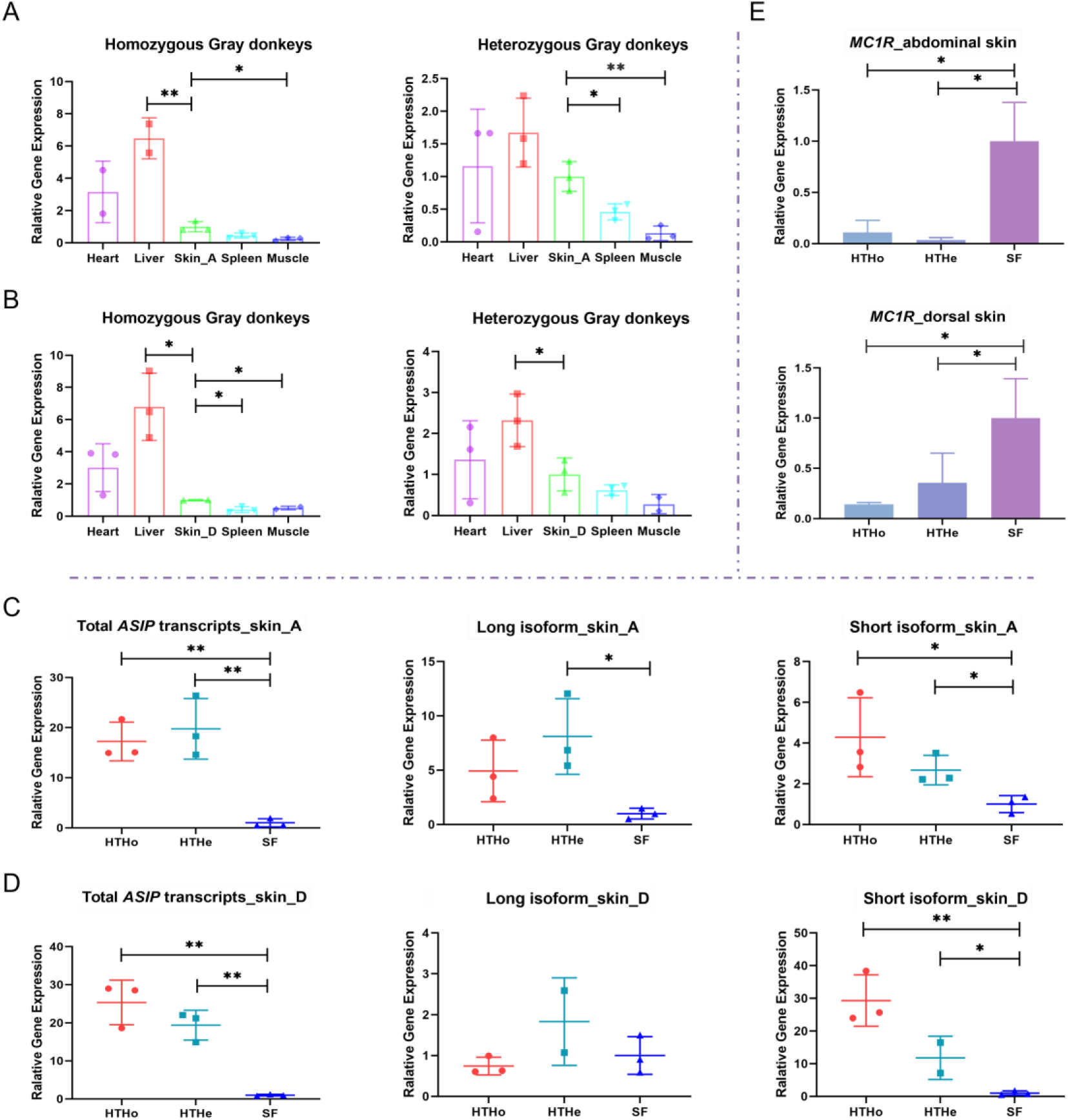
The expression of *ASIP* and *MC1R* in Gray and non-Gray donkeys. **A**: Relative expression levels of *ASIP* in heart, liver, abdominal skin (Skin_A), spleen, and muscle tissues of homozygous Gray donkeys (HTHo) and heterozygous Gray donkeys (HTHe). **B**: Relative expression levels of *ASIP* in heart, liver, dorsal skin (Skin_D), spleen, and muscle tissues of HTHo and HTHe. **C**: Relative expression levels of total *ASIP* transcripts (including both long and short isoforms) in abdominal skin of HTHo, HTHe, and Sanfen individuals of the Dezhou donkey (SF). **D**: Relative expression levels of total *ASIP* transcripts (including long and short isoforms) in dorsal skin of HTHo, HTHe, and SF. **E**: Relative expression levels of *MC1R* in abdominal and dorsal skin tissues of HTHo, HTHe, and SF determined by qPCR. Data are presented as mean ± standard deviation (SD) (n = 3). Statistical significance is indicated as p<0.05 (*) and p<0.01 (**). Skin_A indicates abdominal skin; Skin_D, dorsal skin; HTHo, homozygous Gray donkeys; HTHe, heterozygous Gray donkeys; SF, Sanfen individuals of Dezhou donkeys.

Given that *ASIP* produces both long and short transcript isoforms, transcript-specific qPCR analyses were conducted on dorsal and abdominal skin tissues of homozygous Gray donkeys, heterozygous Gray donkeys, and Sanfen individuals of the Dezhou donkey. In both dorsal and abdominal skin, the total *ASIP* expression measured with the sequences shared by long and short transcripts was significantly higher in homozygous and heterozygous Gray donkeys than in non-Gray donkeys, indicating an overall upregulation of *ASIP* in Gray donkey skin (Fig. 4C, Fig. 4D). Isoform-specific analyses further revealed that the long transcript in dorsal skin did not differ significantly between Gray and non-Gray donkeys regardless of genotypes at *ASIP_*IN2, whereas it was relatively elevated in abdominal skin of heterozygous Gray donkeys (P<0.05). The short transcript in both dorsal and abdominal skin was markedly higher in homozygous and heterozygous Gray donkeys compared with non-Gray donkeys (P<0.05), and showed the highest expression in homozygous Gray donkeys. These results indicate that *ASIP* expression in donkey skin is not only genotype-dependent but also exhibits skin region-specific patterns, and imply the dose effect of G allele at *ASIP*_IN2 on the expression of the short transcript. In addition, qPCR analyses showed that the canonical pigmentation gene *MC1R*, negatively regulated by *ASIP*, had a significant lower expression in Gray donkeys (P<0.05) (Fig. 4E).

## Discussion

In this study, we integrated genome-wide selection scans, fine-scale genotyping, and transcript analyses to elucidate the genetic basis of gray coat color in donkeys. A strong and consistent selection signal was identified on chromosome 15, and multiple evidences converge on a *cis*-regulatory variant within the *ASIP* gene as the major determinant of the gray coat phenotype in Hetian Gray donkeys. By combining population genomic analyses with expression profiling and targeted validation, our results provide the first clear genetic explanation for gray coat color in donkeys and highlight the critical role of regulatory variation in shaping pigmentation traits during animal domestication.

*ASIP* was firstly identified in mouse, and it encodes a paracrine signaling factor and regulates coat color by antagonizing α-melanocyte-stimulating hormone (α-MSH) via binding to *MC1R*, and thereby inhibits eumelanin synthesis and promoting pheomelanin production [36, 37]. Classic mouse studies demonstrated that structural mutations or ectopic expression of *ASIP* could produce agouti or lethal yellow phenotypes, and thus established its central role in melanogenesis [36, 38]. Beyond pigmentation, *ASIP* variants have been found to be involved in diverse physiological processes, including energy metabolism, obesity, embryonic development, and tumorigenesis [39].

In domestic animals, both coding and *cis*-regulatory variants of *ASIP* have been repeatedly shown to contribute to coat color diversity. In dogs, sheep, water buffalo, and rabbits, *ASIP* modulates the ratio of eumelanin to pheomelanin via *MC1R* signaling, often through *cis*-regulatory rewiring mediated by transposable elements [40–45]. In camels, deletions or nonsynonymous mutations in *ASIP* result in darker pigmentation [46–48]. In goats, sheep, alpacas, and camels, *ASIP* expression correlates closely with the development of white, dark, or black coats [3, 49, 50]. In horses, deletions, base substitutions, or 3’UTR variants in *ASIP* can markedly alter protein function or expression levels, and thereby modulate eumelanin production and coat color [46–48, 51], and the interactions between *ASIP* and *MC1R* determine bay and black coat colors in horse. Although studies on coat color in donkeys remain limited, existing evidence indicates that the *ASIP* gene plays a key role in determining donkey coat colors. Black-coated donkeys typically carry a recessive missense mutation (c.349 T>C), resulting in a cysteine-to-arginine substitution at amino acid 117 of ASIP, which affects eumelanin synthesis [9] and may also influence light-point pigmentation [11]. In addition, a 30-bp deletion in *ASIP* was reported to be associated with gray coat color [13]. Together, these cross-species findings indicate that *ASIP* is a recurrent target of selection shaping coat color diversity in mammals. Our study demonstrates that donkey gray coat phenotype is primarily driven by *cis*-regulatory variation in *ASIP*, and reveals its critical role in age-dependent progressive hair depigmentation in donkey.

In the evolutionary context, it is important to examine coat color variation at the population level in Chinese domestic donkeys. Long-term domestication, geographic isolation, and region-specific breeding have shaped substantial phenotypic diversity of donkeys, including black, dun, gray, and spotted coat colors [14]. Notably, smaller-bodied donkeys tend to exhibit dun coloration, whereas larger-bodied donkeys predominantly display black coat with white spots and solid black, and other coat colors are also occasionally observed in the indigenous donkey populations. Despite this diversity, gray coat color remains relatively rare nationwide and exhibits a pronounced localized distribution. The Hetian Gray donkey represents the only indigenous Chinese breed, in which gray coat color has become a stable and defining breed characteristic. In this breed, hair gradually depigments with aging while skin pigmentation remains dark, producing its distinctive gray appearance. Gray coat color is also observed sporadically in other donkey breeds [14]. Consistent with this population pattern, our genetic analyses indicate that the Gray-associated allele is exclusively harbored by the studied Hetian Gray donkeys, whereas it occurs only sporadically in other populations.

Domestic donkeys originated in Africa, and gradually spread across Eurasia [52]. Our results revealed that African wild ass carried the genotype of non-Gray phenotype (Fig. S1), which implies that the gray coat color may appear after donkey domestication. And the fact that Gray donkeys from different breeds carry the same causative mutation suggests that all Gray donkeys have inherited the mutated G allele from a common ancestor. The Xinjiang region was the earliest and critical center of distribution and dispersal of donkeys in China in the ancient time [53]. The Xinjiang donkey, an ancient indigenous breed comprising a few of Gray individuals, has a distribution overlapping that of the Hetian Gray donkey. The Hetian region, located along the southern corridor of the ancient Silk Road, historically functioned as an important hub for trading, and thus facilitated the movement of people and livestock. Our previous study showed that the Hetian Gray donkey may originate from Xinjiang donkeys and Guanzhong donkeys [53]. Taken together, our results suggests that the Gray-associated regulatory variant most likely originated from Xinjiang donkeys or geographically adjacent ancestral populations, rather than arising as a recent de novo mutation unique to the Hetian Gray breed.

Domestic donkeys originated in Africa and subsequently dispersed across Eurasia. Our results showed that African wild asses carried only the non-gray genotype (Fig. S1), suggesting that the gray coat color likely emerged after domestication. Notably, Gray donkeys from different breeds shared the same causative mutation, indicating that the mutated G allele was inherited from a common ancestor rather than arising independently in multiple lineages. The Xinjiang region represented one of the earliest and most important centers of donkey distribution and dispersal in ancient China. The Xinjiang donkey, an indigenous ancient breed with a small proportion of Gray individuals, overlaps geographically with the Hetian Gray donkey. Moreover, the Hetian region, located along the southern route of the ancient Silk Road, historically functioned as a major trade corridor that facilitated gene flow among livestock populations. Our previous study further suggested that the Hetian Gray donkey may have originated from admixture between Xinjiang donkeys and Guanzhong donkeys.

To further investigate the origin of the gray-associated variant, we constructed phylogenetic trees using 8-kb regions flanking the candidate mutation sites (Fig. S2). The results demonstrated that Hetian Gray donkeys clustered more closely with Xinjiang and Guanzhong donkeys. Although the two wild ass species (*Somali wild ass* and *Kiang*) were phylogenetically closest to Xinjiang donkeys, they did not carry the gray-associated allele, thereby excluding them as a direct source of the mutation. These findings support the hypothesis that the gray-associated variant originated within Xinjiang donkey populations or geographically adjacent ancestral groups. Taken together, our results suggest that the gray coat color most likely arose from standing genetic variation in Xinjiang donkeys and was subsequently enriched and maintained through artificial selection, rather than resulting from a recent de novo mutation specific to the Hetian Gray breed.

At the transcript level, our results indicate that the Gray-associated *cis*-regulatory variant is correlated with elevated *ASIP* expression and reduced *MC1R* expression in skin tissue. Notably, this variant is located within intron 2, approximately 4 kb upstream of the initiation of the *ASIP* short isoform, consistent with a potential role in transcript-specific regulation. Gene expression analyses further demonstrated that the short *ASIP* isoform is significantly upregulated and the Gray specific G allele has dose effect on its expression in the skin of Gray donkeys. These findings suggest that enhanced *ASIP* expression may interfere with *MC1R*-mediated melanocyte signaling and potentially affect melanocyte differentiation or survival, and thereby contribute to the progressive hair depigmentation observed in Gray donkeys. Unlike coding mutations that directly alter protein function, this regulatory mechanism allows fine-tuned modulation of gene expression, congruent with the observed age-dependent and spatially variable pigmentation phenotype. Such regulatory variation provides a flexible substrate for selection, enabling phenotypic diversification without compromising essential biological functions.

Notably, hair depigmentation in both horses and donkeys is fully dominant, and both species exhibit age-dependent progressive graying. However, the underlying genetic and cellular mechanisms differ fundamentally between them. In horses, gray coat color is caused by a 4.6-kb duplication within *STX17*, which is associated with abnormal melanocyte proliferation and a high incidence of melanoma [5]. At the molecular pathway level, the *STX17* duplication is thought to influence melanocyte cell-cycle regulation and proliferation-related signaling pathways, potentially involving MAPK/ERK and other networks linked to cell proliferation and tumorigenesis [54]. This results in an abnormal increase in melanocyte number and progressive pigmentary changes. Mechanistically, this represents a dysregulation of melanocyte proliferation leading to the gray phenotype. In contrast, gray coat color in donkeys is mediated by a *cis*-regulatory variant in *ASIP*, which does not involve known melanoma-associated pathways, and no melanoma has been observed in Gray donkeys. *ASIP* encodes the Agouti signaling protein, which antagonizes the MC1R signaling pathway, suppressing eumelanin synthesis while promoting pheomelanin production. Through this mechanism, *ASIP* regulates the relative proportion of melanin types and alters pigment expression patterns [55]. Unlike the mechanism in horses, this pathway primarily affects melanin synthesis rather than melanocyte number, and therefore does not involve abnormal cellular proliferation or tumorigenesis. Gray coat color in donkeys can be further divided into several subtypes, suggesting the involvement of potential modifying genes. Several limitations of this study should be acknowledged. First, coat color phenotypes among donkey breeds can be visually similar, particularly among light or mixed-color individuals, which may introduce ambiguity in phenotype classification. Second, incomplete historical records for some individuals limited our ability to precisely trace phenotype–genotype relationships across generations. Future studies incorporating longitudinal phenotypic documentation, functional assays, and larger population datasets will be essential for further elucidating the regulatory mechanisms by which *ASIP* influences age-related coat color phenotype in donkeys.

## Conclusions

In summary, we identified a *cis*-regulatory variant in *ASIP* as the key genetic determinant of the gray coat color in donkeys and demonstrated that similar age-related depigmentation phenotypes in equids can arise through distinct genetic pathways. These findings advance our understanding of coat color genetics in donkeys and provide broader insight into the evolutionary and regulatory mechanisms underlying phenotypic diversity in domesticated animals.

## Supplementary Information

**Supplementary Material 1: Fig. S1** Evolutionary conservation of the candidate mutation site on *ASIP* across mammals. Multiple sequence alignment of the genomic region encompassing the candidate SNP across *Equus asinus*, *African wild ass*, *Equus caballus*, *Sus scrofa*, *Felis catus*, *Homo sapiens*, *Ovis aries*, *Macaca mulatta* and *Bos taurus*, highlighting the evolutionary conservation of the surrounding sequence and the candidate nucleotide position (*ASIP*_IN2).

**Supplementary Material 2: Fig. S2** Phylogenetic analysis based on 8-kb flanking sequences surrounding the gray-associated regulatory variant (HT: Hetian Gray donkey; GZ: Guanzhong donkey; GL: Guangling donkey; XJ: Xinjiang donkey; Somali: *Equus africanus somaliensis*; Kiang: *Equus Kiang*)

**Supplementary Material 3: Table S1** Primers and PCR conditions for amplifying and sequencing the variant in intron 2 of *ASIP*

**Supplementary Material 4: Table S2** Mass spectrometry–based genotyping results of Gray and non-Gray donkeys

**Supplementary Material 5: Table S3** Single Nucleotide Polymorphisms Identified Within Candidate Regions

## Acknowledgments

We acknowledge the High-Performance Computing Platform of China Agricultural University for providing computational support.

## Author contributions

YYL conducted the experiments and data analysis, and prepared the manuscript. YYL and YL performed the study design. JW, SL, XL, and KG contributed to data collection. TY and MF contributed to the statistical analysis. YYL, HZ, XW, WX, SQ, RY and CZ contributed to interpreting results and critically revising the manuscript. YYL, YL, TY, and MF performed the collection and preparation of DNA samples. CZ conceived and coordinated the study, and revised the manuscript. All authors read, revised, and approved the final manuscript.

## Data availability

The genome resequencing data for the Xinjiang donkey are available upon request.

## Declarations

### Ethics approval and consent to participate

The samples were obtained following the principle approved by the Animal Care and Use Committee of China Agricultural University (permit number: XK257). The informed consent from the owners were obtained for using the samples of the donkeys in the present study.

### Consent for publication

Not applicable

